# Change in the bladder function of rats with focal cerebral infarction induced by photochemically-induced thrombosis

**DOI:** 10.1101/2021.08.04.455101

**Authors:** Yuya Ota, Yasue Kubota, Yuji Hotta, Mami Matsumoto, Nayuka Matsuyama, Taiki Kato, Takashi Hamakawa, Tomoya Kataoka, Kazunori Kimura, Kazunobu Sawamoto, Takahiro Yasui

## Abstract

The photochemically-induced thrombosis(photothrombosis) method can create focal cerebral infarcts anywhere in the relatively superficial layers of the cerebrum; it is easy to implement and minimally invasive. Taking advantage of this versatility, we aimed to establish a new rat model of urinary frequency with focal cerebral infarction, which was characterized by its simplicity, nonlethal nature, and high reproducibility. The prefrontal cortex and the anterior cingulate cortex, which are urinary centers, were targeted for focal cerebral infarction, and urinary parameters were measured by cystometrogram. Cystometric analysis indicated that micturition intervals significantly shortened in photothrombosis-treated rats compared with those in the sham operative group on Days 1 and 7 (P < 0.01), but prolonged after 14 days, with no difference between the two groups. Immunopathological evaluation showed an accumulation of activated microglia, followed by an increase in reactive astrocytes at the peri-infarct zone after photothrombotic stroke. Throughout this study, all postphotothrombosis rats showed cerebral infarction in the prefrontal cortex and anterior cingulate cortex; there were no cases of rats with fatal cerebral infarction. This model corresponded to the clinical presentation, in that the micturition status changed after stroke. In conclusion, this novel model combining nonlethality and high reproducibility may be a suitable model of urinary frequency after focal cerebral infarction.

## INTRODUCTION

Cerebral infarction is associated with a high incidence of lower urinary tract symptoms (LUTS), particularly urinary bladder overactivity, which reduces a patient’s quality of life [1,2]. Therapies that target bladder receptors, such as anticholinergics and beta-3 adrenaline receptor stimulants, have been used to treat LUTS caused by stroke, but they are not sufficiently effective. Currently, there is no established LUTS treatment targeting the central nervous system. We propose that one of the reasons for the lack of progress in treatments for the central nervous system is the dearth of animal models.

Middle cerebral artery occlusion (MCAO) rats have generally been used as a model for bladder overactivity caused by cerebral infarction. However, they have the disadvantages of variability in the infarct volume and a high mortality rate due to the large infarct area and local traumatic effect [3,4,5]. Furthermore, creating an MCAO rat model is surgically demanding, and it takes a long time before such a model can be produced consistently [6].

Establishing a simple and reproducible animal model of LUTS caused by cerebral infarction is needed to solve these issues. Therefore, we focused on applying photochemically-induced thrombosis (photothrombosis) as a method for creating focal cerebral infarcts [7]. The principle of arterial occlusion achieved by photothrombosis is the formation of a stable thrombus that consists solely of aggregated platelets in response to endothelial peroxidative damage in laser irradiated areas [8]. Depending on where the light is applied, photothrombotic reaction can create focal cerebral infarcts anywhere in the relatively superficial layers of the cerebrum. Although photothrombosis has been used in various brain experiments [9,10], it has never been applied for studying urinary function.

The periaqueductal gray (PAG) region receives sacral afferents related to bladder filling and transmits efferent signals to the sacral spinal cord or bladder via the pontine micturition center (PMC) [11]. Previous reports have shown that the prefrontal cortex (PFC) and the anterior cingulate cortex (ACC) exert supra-inhibitory control of these voiding reflexes both experimentally and clinically [12,13,14,15], indicating that urination can be initiated and inhibited voluntarily by PFC and ACC. In other words, a cerebral infarction of PFC and ACC may impair urinary control and cause subsequent urinary bladder overactivity. There are few reports of histopathological observation of the phenomena occurring in the brain in a model of bladder overactivity caused by cerebral infarction, although various experiments related to the mechanism of urination have been conducted [13,16].

Our aim was to establish a new rat model of urinary frequency, with focal cerebral infarction in PFC and ACC induced by photothrombosis, which is easy to perform and minimally invasive, and to observe histopathological changes in the brain.

## MATERIALS AND METHODS

### Experimental design and animals

All experimental protocols were approved by the Animal Care Committee of Nagoya City University Graduate School of Medical Sciences, Nagoya, Japan (No. H30M-57). The study design is shown in Fig. 1. A total of 90 11-week-old female Wistar-ST rats were purchased from SLC Inc. (Shizuoka, Japan). Ten rats were allocated for the evaluation of the infarct area and volume, in a small pilot study, and 40 rats each were assigned to a sham group and a photothrombosis-treated group. Neurological evaluation, cystometric measurements, and immunohistochemical evaluation of the brain were performed preoperatively and on postoperative Days 1, 7, 14, and 28, (n = 8). We did not correct cystometrogram data for some rats because of cannula withdrawal during measurement. Thus, the final preoperative numbers were: n = 8 (Day 1, n = 8; Day 7, n = 6; Day 14, n = 7; Day 28, n = 7) in the sham group, and n = 8 (Day 1, n = 7; Day 7, n = 8; Day 14, n = 7; Day 28, n = 8) in the photothrombosis group. The rats were maintained at 21 ± 2ºC with a 12-h light-dark cycle and were allowed free access to water and provided a standard laboratory diet.

**Figure 1.**
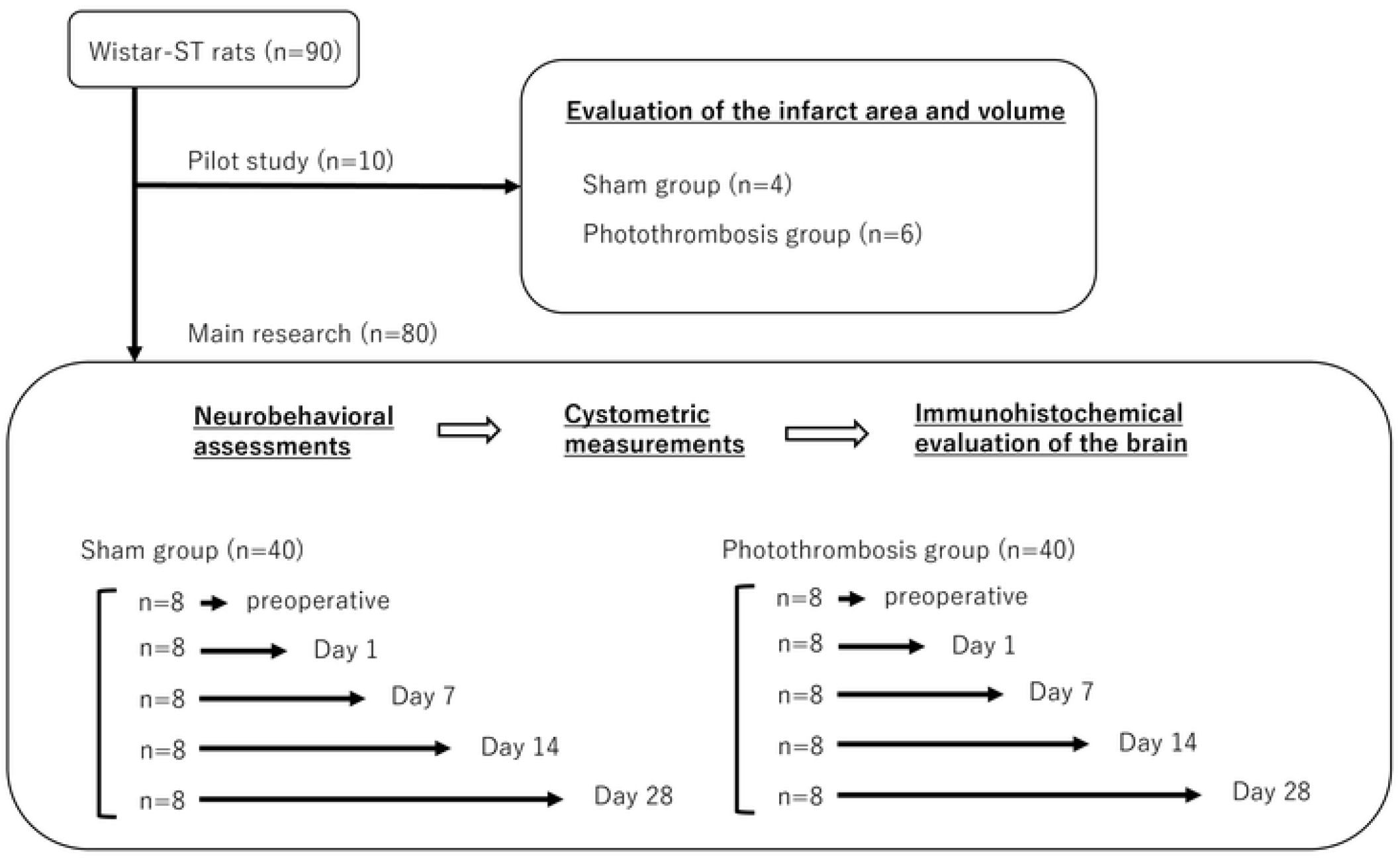
Study design

### Photothrombosis

We used a modified version of the photothrombosis technique described by Watson to induce cerebral infarction [7]. The rats were subjected to 4% isoflurane inhalation for induction and 1.5%–2.0% isoflurane for maintenance. The scalp was incised to expose the skull surface and cleaned to reveal bregma and the target area for irradiation (Fig. S1A). For illumination, a fiber optic cable delivering a light source of 8 mm in diameter (MSG6-1100S, Moritex, Saitama, Japan) was placed stereotactically onto the skull 1 mm anterior to the bregma on the midline (Fig. S1B). Cerebral infarction was induced by activation of photosensitive Rose Bengal dye (30 mg/kg, 330000-1G; Sigma-Aldrich, St. Louis, MO, USA) in 0.9 % NaCl solution. Two minutes prior to laser irradiation, Rose Bengal was injected through the caudal vein. The skull was illuminated for 30 minutes (wavelength 533 nm; 150 mW, MHAA-100W-100V, Moritex). Sham-operated rats received normal saline instead of Rose Bengal solution. After laser irradiation, the scalps were sutured, and the animals were allowed to recover in their home cages.

### Assessment of infarction aria and volume

It is known that 2,3,5-triphenyl tetrazolium chloride (TTC) (T8877, Sigma-Aldrich) staining allows for rapid assessment of the infarcted area [17]. This TTC staining can be used only in the first few days after stroke onset due to the action of mitochondria1 enzyme systems [18]. Therefore, 1 day after photothrombosis, TTC staining was used to assess the area of cerebral infarction. The rats were euthanized in a chamber filled with carbon dioxide. Following sacrifice, the brains were removed from the skull and stored at −20°C for 10 min. The area of PFC and ACC is located 0.4 mm lateral to the midline in a sagittal section, according to the stereotactic section map designed by C. Watson and G. Paxinos [19]. Hence, the cerebral hemispheres were cut into sagittal slices, each 2-mm thick, based on a distance of 0.4 mm from the midline, and incubated in 4% TTC–PBS at 37°C for 30 min. The area of infarction in each slice was measured using a digital scanner and Image J software (version 1.52u; National Institutes of Health, Bethesda, MD, USA). The volume of infarction in each animal was obtained as the product of average slice thickness and sum of infarction areas in all brain slices examined (infarct volume = area of infarct [mm2] × thickness [2 mm]).

### Neurological evaluation

The modified neurological severity score (mNSS) was used to evaluate neurobehavioral functions [20]. The mNSS includes motor, sensory, reflex, and balance tests. Neurological function was graded on a scale of 0–18 (normal score, 0; maximal deficit score, 18). For severity, points are scored for the inability to perform a test or the lack of examined reflexes; thus, the higher the score, the greater the severity of the injury.

### Cystometric investigations

After the neurological evaluation, all rats received cystometry according to methods described previously [21,22,23]. With inhalation anesthesia (4% isoflurane for induction and 1.5%–2.0% isoflurane for maintenance), the abdomens of the rats were opened through a midline incision, and the bladders were exposed. A polyethylene catheter tip (PE-50: inner diameter 0.5 mm, outer diameter 1.0 mm; Eastsidemed, Tokyo, Japan) was heated to make a collar that could fit tightly into the bladder. It was inserted into the bladder dome and sealed by a purse-string suture (5-0 silk suture) under a surgical microscope. After suturing, the bladder was filled with saline solution up to the leak point and a leak at the urethral meatus was observed, confirming that no leak had occurred at the suture site. The other end of the catheter was tunneled subcutaneously to the neck and connected to a syringe pump (U-802, Univentor, Zejtun, Malta) that was attached to a pressure transducer (MLT844; ADInstruments, Dunedin, New Zealand) to allow saline infusion. After suturing the abdominal skin, the rats were placed in a restraining cage (ICN-5; Tokyo Garasu Kikai, Tokyo, Japan) for the cystometrogram. Three hours after awakening from anesthesia, the bladders were filled with saline (room temperature) at 0.10 mL/min for 120 min, and intercontraction intervals (ICIs), baseline pressure (BP), micturition threshold pressure (TP, bladder pressure immediately prior to micturition), and maximum intravesical pressure (MP) were recorded for analysis. After the cystometrogram, the entire bladder was collected and weighed.

### Brain preparation for histology

We used NeuN (1:500; mouse monoclonal, MAB377, Sigma-Aldrich) as a neural marker to assess the distribution of mature neurons. The area from which mature neurons had disappeared indicated the infarcted area. Anti-ionized calcium binding adaptor molecule 1 (Iba1) (1:2000; Rabbit polyclonal, 013-27691, Wako, Osaka, Japan) and anti-glial fibrillary acidic protein (GFAP) (1:2,000; mouse monoclonal, G3893, Sigma-Aldrich) were used to assess activated microglia and reactive astrocytes in ischemic regions, respectively. After cystometrogram, the rats were exsanguinated from their right auricle, following inhalant anesthesia using isoflurane, and transcardially perfused with 20 mL/body of 1% heparinized saline followed by 25 mL/body 10% formalin. The brains were removed, postfixed overnight for 24 additional hours in 10% formalin at 4°C, and embedded in paraffin. All brain tissue were sectioned into 4-μm-thick sagittal cross-sections of PFC and ACC, which is located 0.4 mm lateral to the midline, as described above. The sections were deparaffinized in D-limonene and 100% ethanol (3 times for 10 minutes) before rehydration in graded ethanol (90%, 80%, and 50%; 2 minutes each), followed by MilliQ-H_2_O (5 minutes). Antigen retrieval was performed by heating the sections in 10 mM trisodium citrate dehydrate (pH 6) in Decloaking Chamber^™^ NxGen (Biocare Medical, Pacheco, CA, USA) at 95°C for 40 minutes before cooling at room temperature for 30 minutes. After blocking the endogenous peroxidase activity by methanol containing 3% H_2_O_2_, the brain slice tissues were incubated overnight at room temperature with antibody against NeuN. After washing in buffer, the sections were immunostained by the avidin–biotin peroxidase method using the Vectastain Elite Kit (PK-6102, Vector Laboratories, Burlingame, CA, USA) with 3–3-diaminobenzidine and hydrogen peroxide as the chromogen. For immunofluorescence staining, the brain slice tissues were incubated with anti-Iba1 and anti-GFAP overnight at 4°C, and incubated with goat anti-rabbit or goat anti-mouse secondary antibodies (1:2000; Zhongshan Goldbridge Biotechnology, Beijing, China) for 1 h at 25°C. For immunostaining and immunofluorescence analyses, BZ-X800 (Keyence, Osaka, Japan) was used to acquire digital images which were analyzed with a BZ-X800 Analyzer.

For quantification of immunofluorescence in the ischemic region, which is within 300 μm beneath the infarcted area, as indicated by NeuN staining, five microscopic images per rat were randomly captured at 40× objective using the microscope, according to previous reports [24]. In the sham group and preoperatively in the PT group, the images were taken randomly within the frontal lobe. The numbers of Iba1-, and GFAP-positive cells with identified same-threshold nuclei (4′,6-diamidino-2-phenylindole [DAPI]-stained) were measured with a BZ-X800 Analyzer.

### Statistics

All statistical analyses were performed using the Statistical Package of Social Sciences 22 software® (SPSS; Chicago, IL, USA). The results are expressed as mean ± standard deviation (SD). Student’s t-tests were used for comparison between the two groups. A two-way repeated measures analysis of variance (ANOVA) was used to perform multiple group comparisons, followed by Tukey’s test for individual comparisons. The results were considered statistically significant at P < 0.05.

## RESULTS

### Evaluation of infarction area and volume

As shown in Fig. 2, the sagittal brain sections on Day 1 after photothrombosis revealed infarcted PFC and ACC lesions. All the rats in this study had similar areas of infarction (Fig. S2). No damage was observed in the brain stem, such as PAG or PMC regions, in either group. The mean infarction volume was 147.1 ± 24.1 mm3. No evidence of infarction was found in any of the sham rats.

**Figure 2.**
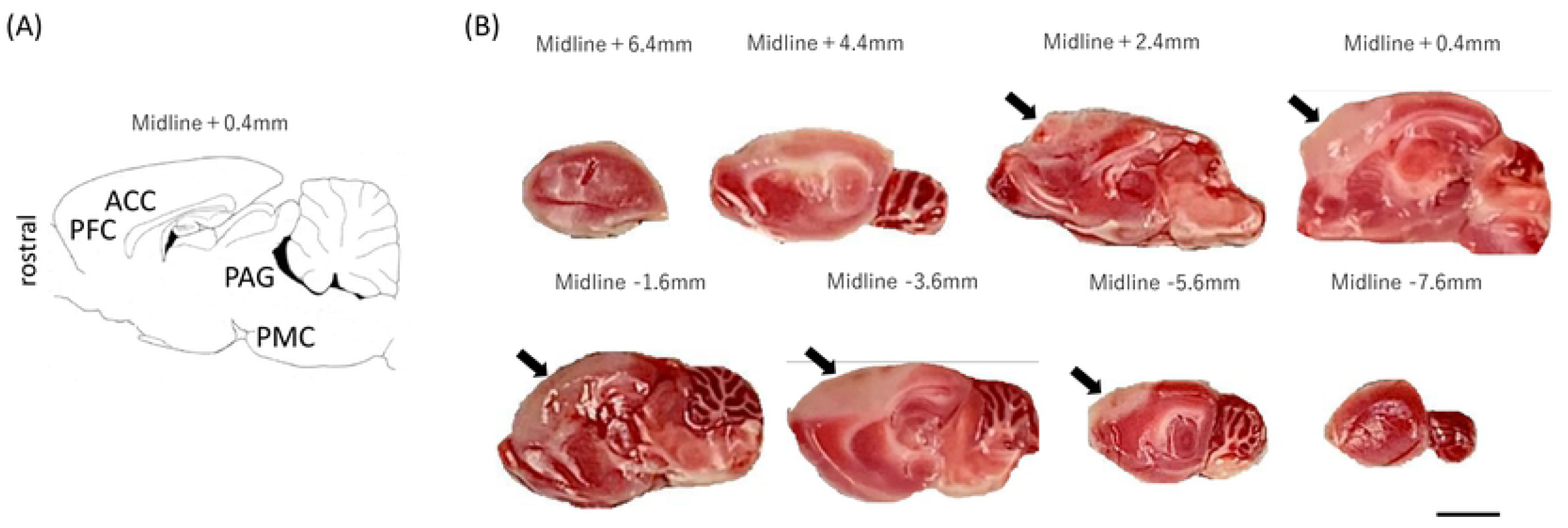
Diagrammatic representation of a sagittal section and triphenyl tetrazolium chloride (TTC)-stained brain images A: Anterior cingulate cortex (ACC), prefrontal cortex (PFC), periaqueductal gray (PAG), pontine micturition center (PMC) are present in the sagittal section 0.4 mm lateral to the midline identified according to the rat brain atlas of Paxinos and Watson. B: Examples of TTC-stained sagittal sections of rat brains on Day 1 after photothrombosis. Cerebral infarction was identified in PFC and ACC in the brains of all rats after photothrombosis. Arrows indicate unstained areas (areas of cerebral infarction). The distances from the midline are shown above each brain slice. Scale bars = 3 mm.

### General information

The body and bladder weights are shown in Table 1. There were no cases of rats with fatal cerebral infarction during the postoperative period. There were no statistical differences in body weight, bladder weight, or bladder body weight ratio (bladder weight/body weight) between the sham group and the photothrombosis group (PT group) at any time points.

**Table 1.** Body and bladder weights in the different groups.

|  | Sham group | Photothrombosis group | P-value |
| --- | --- | --- | --- |
| Body weight, g |  |  |  |
| Preoperative | 218.4 ± 7.3 | 219.3 ± 3.0 | 0.758 |
| Day 1 | 214.9 ± 10.2 | 209.9 ± 6.2 | 0.208 |
| Day 7 | 227.8 ± 10.1 | 221.9 ± 10.7 | 0.312 |
| Day 14 | 242.4 ± 8.5 | 238.1 ± 6.3 | 0.303 |
| Day 28 | 257.3 ± 4.2 | 252.4 ± 7.2 | 0.138 |
| Bladder weight, mg |  |  |  |
| Preoperative | 151.6 ± 17.8 | 146.5 ± 13.8 | 0.531 |
| Day 1 | 156.9 ± 20.7 | 154.1 ± 21.9 | 0.808 |
| Day 7 | 164.0 ± 13.9 | 156.9 ± 19.3 | 0.459 |
| Day 14 | 160.9 ± 17.4 | 163.4 ± 15.8 | 0.777 |
| Day 28 | 166.1 ± 18.9 | 170.9 ± 20.0 | 0.647 |
| Bladder body weight ratio<br>(×10<sup>-3</sup>) |  |  |  |
| Preoperative | 0.69 ± 0.08 | 0.67 ± 0.06 | 0.452 |
| Day 1 | 0.73 ± 0.09 | 0.73 ± 0.10 | 0.919 |
| Day 7 | 0.72 ± 0.05 | 0.71 ± 0.10 | 0.818 |
| Day 14 | 0.66 ± 0.08 | 0.69 ± 0.07 | 0.599 |
| Day 28 | 0.65 ± 0.03 | 0.68 ± 0.02 | 0.421 |

### Modified neurological severity score

Figure 3 show the results of neurobehavioral assessments with the mNSS test for each rat. All of the individual data points are detailed in the Supporting Information (File S1). Immediately after photothrombosis, the rats showed characteristic motor changes, with bilateral hind limbs flexed when suspended by the tail. Furthermore, rats with cerebral infarction from photothrombosis were observed to have balance disorders in the beam balance test (walking on a 3-cm beam); however, there were no sensory or reflex disturbances. There were no abnormal neurological findings in the sham group at all. Therefore, the mNSS was significantly higher in the PT group than in the sham group at all time points (P < 0.05). The neurological deficits tended to improve with time, but neurological damage was still present 28 days after the stroke.

**Figure 3.**
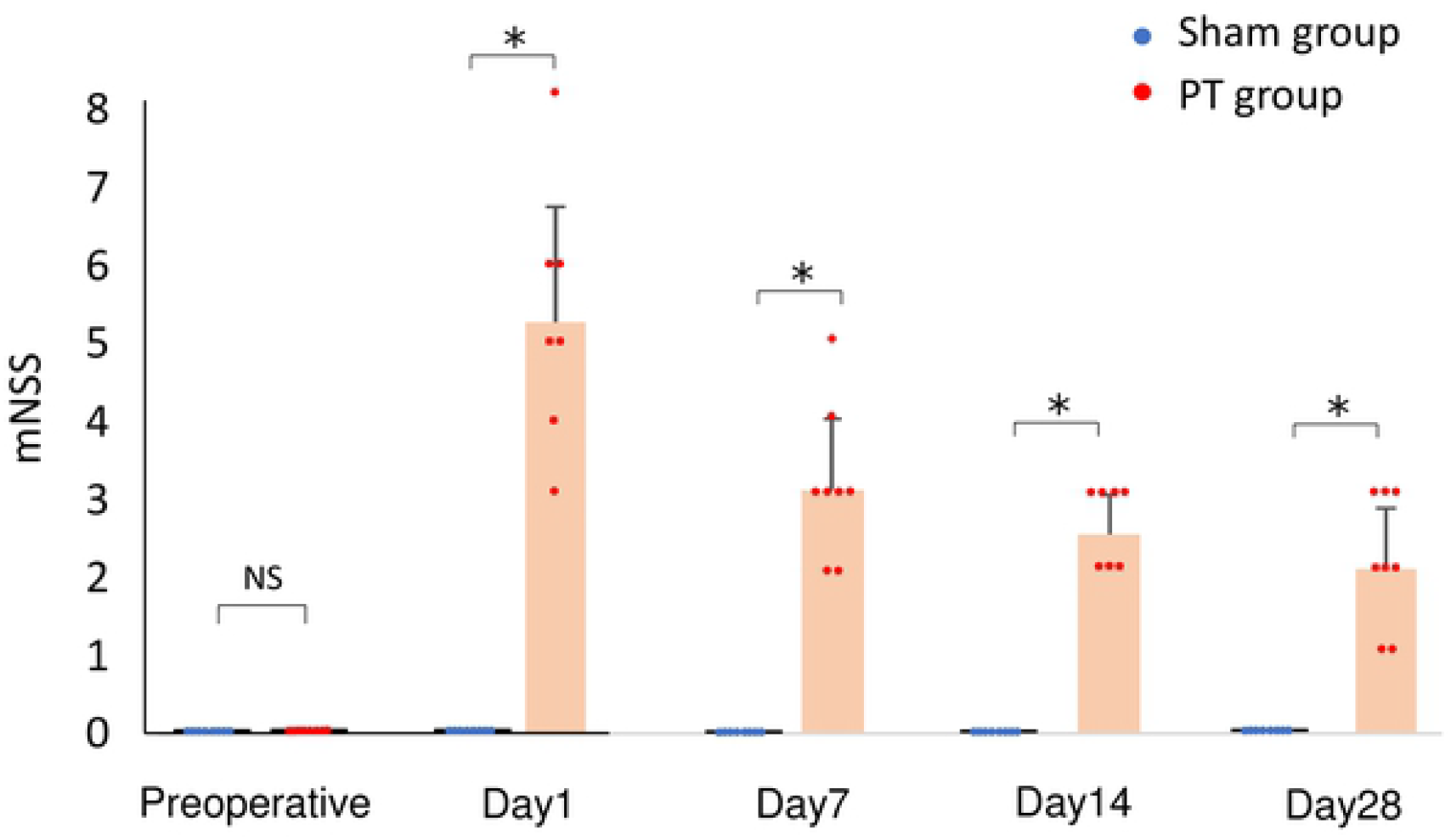
Time course of changes in the modified neurological severity score Neurological functional tests were performed in the sham and photothrombosis (PT) rats at each time point. There were significant differences (*P < 0.001, Student’s t-test) in the mNSS of each group. Values represent mean ± SD.

### Assessment of the peri-infarct area

In the PT group, mature neurons were lost in the frontal lobe of the cerebral cortex and the ACC from Days 1 to 28, whereas they were present in the sham group and preoperatively in the PT group (Fig. 4A). Immunofluorescence staining revealed positive cells for Iba1 and GFAP staining in the peri-infarct zone (Fig. 4B). Quantitative immunofluorescence staining showed an increase in the number of Iba1- and GFAP-positive cells in the PT group compared with the sham group, from Day 1 to Day 28 and from Day 7 to Day 28 after photothrombosis, respectively (Fig. 4C).

**Figure 4.**
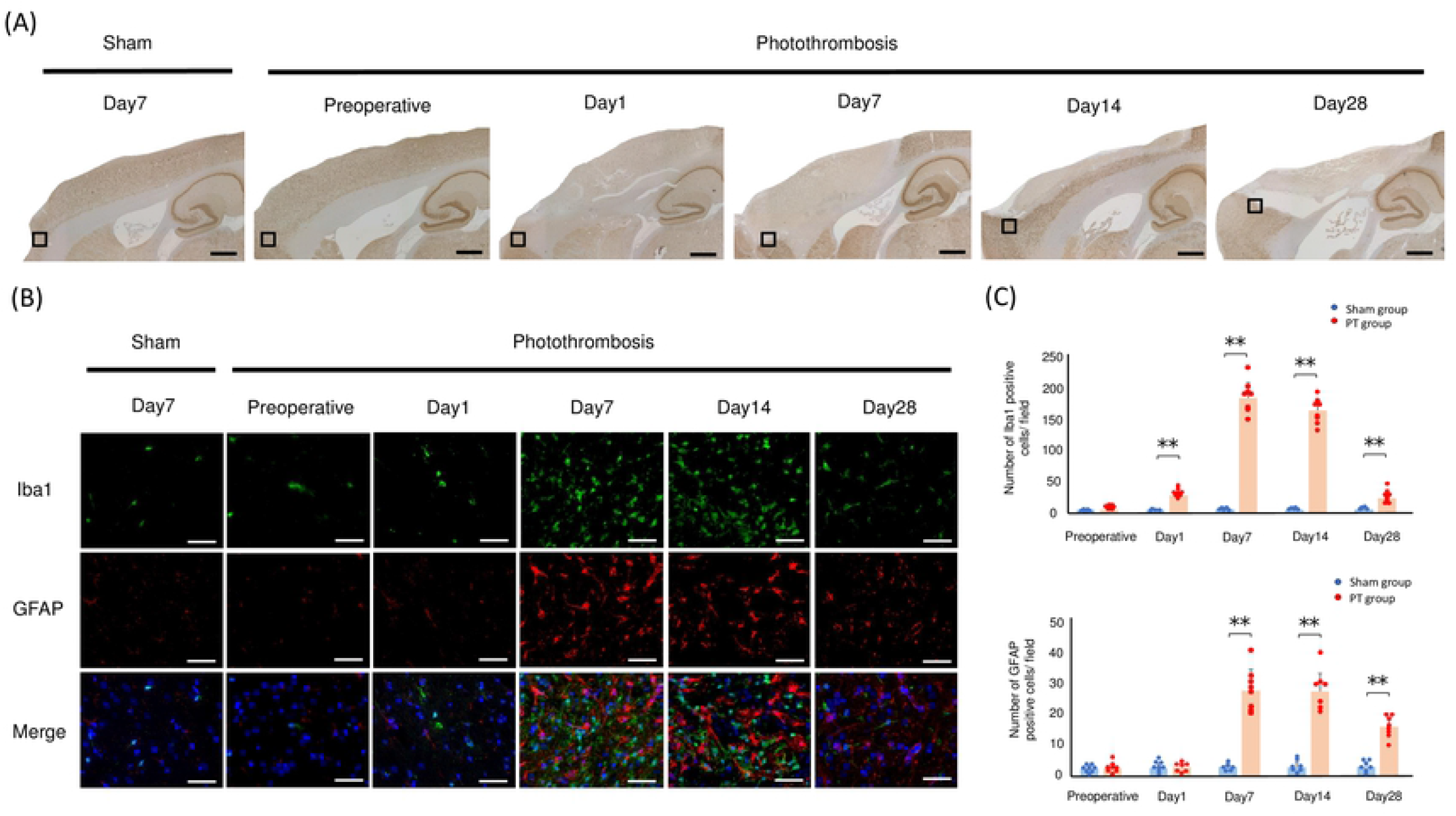
NeuN staining of sagittal sections and sustained activation of glial cells in ischemic regions A: Schematic of a sagittal section (0.4 mm lateral to the midline) centered on the frontal lobe. The black square indicates the location of the peri-infarct zone. Scale bars: 1.5 mm. B: Immunofluorescence staining revealed activated microglia labeled with anti-ionized calcium binding adaptor molecule 1 (Iba1; green), reactive astrocytes labeled with anti-glial fibrillary acidic protein (GFAP; red), and nuclei labeled with 4′,6-diamidino-2-phenylindole (DAPI; blue) in ischemic regions after photothrombosis. Scale bar = 50 μm. C: Quantification of Iba1-positive and GFAP-positive cells. Significant differences between the photothrombosis (PT) group and sham group are indicated by ** (P < 0.001, Student’s t-test). Values represent mean ± SD.

### Cystometric investigations

The representative cystometric curves of each group are shown in Fig. 5A. The cystometric studies showed that ICIs in the PT group were significantly shorter than those in the sham group on Days 1 and 7 (P < 0.01) and ICIs on Day 14 after photothrombosis was significantly longer than those on Day 1 after photothrombosis (P < 0.05). However, there was no significant difference between the sham and photothrombosis groups from Day 14 onwards (Fig. 5B); this means that micturition status in the PT group improved after exacerbation. BP, TP, and MP did not differ between the two groups at any time point (Fig. 5C–E).

**Figure 5.**
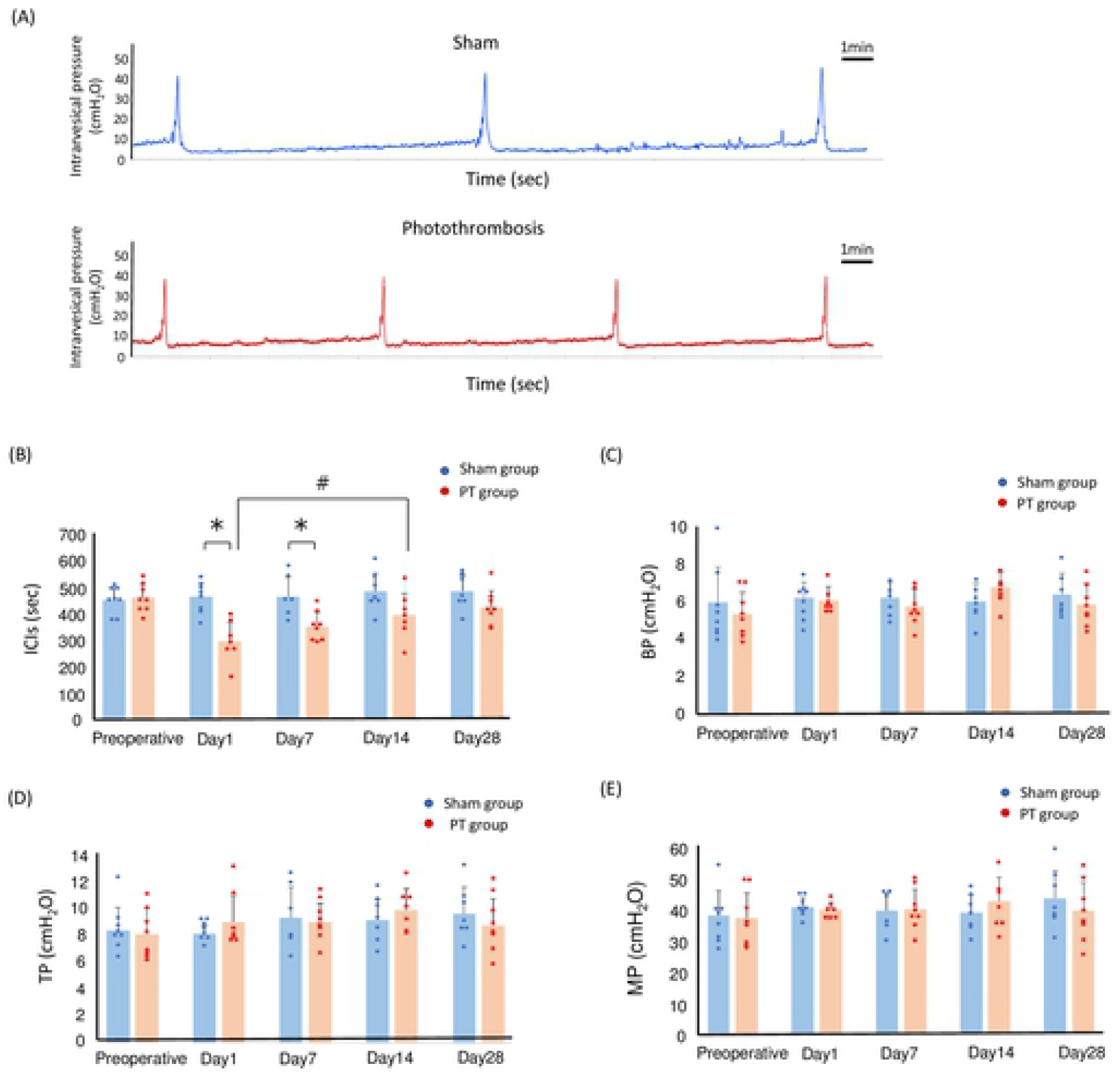
Cystmetric study of the photothrombosis rats compared with the sham rats A: The representative cystometric chart of each group on Day 1 after photothrombosis. B, C, D, and E: Time courses of the changes in intercontraction intervals (ICIs) (B), baseline pressure (BP) (C), micturition threshold pressure (TP, bladder pressure immediately prior to micturition) (D), and maximum intravesical pressure (MP) (E) of the sham versus the photothrombosis (PT) rats. Significant differences between ICIs in the sham and photothrombosis rats are indicated by * (P < 0.05, Student’s t-test). Significant differences between ICIs on Day 1 and Day 14 in the photothrombosis rats are indicated by # (P < 0.05, two-way ANOVA followed by Tukey’s test for individual comparisons). Values represent mean ± SD.

## DISCUSSION

In this study, we used a photothrombotic stroke method to induce focal infarcts in PFC and ACC, and we observed decreased ICIs on Days 1 and 7, according to the cystometric analysis. Furthermore, all rats survived after the operation. To the best of our knowledge, this is the first study in which photothrombosis has been applied to measure urinary function.

The method of inducing a stroke in the cerebral cortex using photochemical reactions was developed by C. Watson in 1985 [7]. Photothrombosis has the advantage of reproducibility due to precise control of the size and location of the infarct, with similar characteristic inflammation and neurodegeneration to other established stroke models [8,25]. Hence, photothrombosis can target urinary centers such as PFC and ACC for stroke lesion. Additional advantages of the photothrombotic model are the minimal surgical invasiveness required and the low mortality rate, as this method does not require opening of the dura [26]. The greater the extent of the infarct, the more likely it is that bladder overactivity will develop, but other disorders will also increase, leading to increased mortality. We have created a model of urinary frequency with only minimal periprocedural dysfunction, without suffering an infarction in the cerebellum, brainstem, or other limbic system, which is necessary to sustain vital functions.

There was no difference in bladder weight or bladder to body weight ratio between the sham and PT groups, which is the same result as obtained in the MCAO model [16]. This suggests that urinary frequency is due to dysfunction of the central nervous system. Yokoyama et al. reported that bladder overactivity after MCAO is the result of upregulation of the NMDA glutamatergic pathway from the forebrain, which promotes a sustained excitatory signal to the PMC [13,27]. Additionally, they state that bladder overactivity after MCAO persisted for 4 months. However, it has been clinically observed that urinary status changes with time in patients with stroke, depending on age, stroke severity, and other disabling diseases [28]. In the present study, rats with cerebral infarction induced by photothrombosis developed urinary frequency for a period of 1 week after ischemia, and showed improvement in urinary function by Day 14 after the onset of stroke. In this respect, we can suggest that our animal model reflects the human clinical presentation. Our model revealed early improvement in urinary frequency, which may be due to the fact that the volume of cerebral infarcts by our method was smaller than that induced by the MCA occlusion method.

Once necrotic, mature neurons do not recover, but the urinary status of the rats changed after stroke in this study. It is possible that the surrounding tissue near the necrotic lesion, which is called the penumbra, compensated for the loss function. Microglia, the immune cells resident in the brain, are the first to respond to ischemic neurons and promote their survival [29]. Following this, it has been suggested that reactive astrocytes in the penumbra contribute to brain remodeling, such as neuronal circuitry and tissue reorganization [30]. Activations of these glial cells within the peri-infarct territory after focal cerebral infarction were reported to occur rapidly within a few days and were sustained for 14 days [31,32]. We suggest that the role of glial cells, such as microglias and astrocytes, has beneficial influence on the early improvement of dysuria in the PT model. Furthermore, complementation of function by structurally undamaged distant areas is a known mechanism of recovery after brain injury [30, 33]. Other micturition centers, such as the insula, hypothalamus, and PAG, which have connections with PFC and ACC, may complement the voiding function [11,34]. Although this study has a limitation in that it is not directly comparable to the MCAO model, we have made the first immunopathological observations of the brain in a model of overactive bladder caused by stroke. We believe that this novel animal model, which reflects changes in voiding function as with clinical symptoms, may significantly contribute to development of new treatments for lower urinary tract dysfunction.

## CONCLUSIONS

Our photothrombosis model was characterized by its simplicity, ability to create localized infarcts, nonlethal nature, and high reproducibility. Similar to the clinical course of a patient after stroke, this model observed changes in micturition status. In other words, this model would be suitable as a new model of urinary frequency caused by cerebral infarction.

## ACKNOWLEDGMENTS

The authors acknowledge the assistance of Naomi Kasuga in the experiments. This study was financially supported by GSK Japan Research Grant 2019 and a grant awarded by the Japan Society for the Promotion of Science, Grant-in-Aid for Young Scientists (B) (Grant Numbers 19k18568) (all to Y.O.).

## AUTHOR CONTRIBUTIONS

Conceptualization: Yuya Ota, Yasue Kubota

Formal analysis: Yuya Ota, Yuji Hotta, Taiki Kato

Funding acquisition : Yuya Ota

Investigation: Yuya Ota, Yuji Hotta, Nayuka Matsuyama, Taiki Kato, Takashi Hamakawa

Methodology: Yuya Ota, Mami Matsumoto, Tomoya Kataoka

Project administration: Yuya Ota, Yuji Hotta

Resources: Nayuka Matsuyama, Taiki Kato, Takashi Hamakawa

Software: Yuya Ota

Supervision: Yasue Kubota, Kazunobu Sawamoto, Kazunori Kimura, Takahiro Yasui

Validation: Yuji Hotta

Writing – original draft: Yuya Ota

Writing – review & editing: Yuji Hotta, Yasue Kubota, Kazunobu Sawamoto, Kazunori Kimura, Takahiro Yasui

## Supporting information

**Figure S1**. Photograph of the creation of photothrombosis

A: Identification of the bregma, the craniometric point at the junction of the sagittal and coronal sutures at the top of the cranium, marked with a black dot indicated by yellow arrow. B: A fiber optic cable delivering a light source was fixed 1 mm anterior to the bregma on the midline under anesthesia inhalation. After Rose Bengal was injected, the skull was illuminated.

**Figure S2**. All triphenyl tetrazolium chloride (TTC)-stained sagittal brain sections at Day 1

2-mm slices of all brains in all sham and photothrombosis rats in the pilot study.

**File S1**. All individual data points in modified neurological severity score The spreadsheet contains the raw data points used to create the graph.

